# Chronic circadian disruption in adolescent mice impairs hippocampal memory disrupting gene expression oscillations

**DOI:** 10.1101/2024.10.18.619106

**Authors:** Ines Gallego-Landin, Paula Berbegal-Sáez, Olga Valverde

## Abstract

Chronodisruption, the misalignment of internal biological systems with external environmental changes, leads to adverse health effects. Particularly, Social Jet Lag (SJL) is defined as the discrepancy between social and biological time and it exemplifies this misalignment, affecting a large part of the young population and impacting cognitive function. Despite its prevalence, our understanding of how developmental chronodisruption ultimately translates into morbidity is limited. To address this, we implemented a chronic chronodisruption protocol in adolescent mice consisting of light/dark cycle manipulation. We employed a comprehensive battery of established behavioral tests alongside an in-depth analysis of the oscillatory expression of the molecular clock and other genes involved in relevant physiological function. Our results show that chronic circadian disruption during adolescence induces impairments in short-term, social, and spatial memory without prompting anxiety-like behavior. Additionally, we report altered gene expression patterns of circadian clock genes *per1*, *per2*, *cry2* and *npas2* in the hypothalamus and the hippocampus. Lastly, we observed a disruption of hippocampal gene expression oscillations which may underlie the hippocampal memory impairments. Overall, this work underscores the critical role of adolescent circadian rhythms in maintaining cognitive function, the relevance of circadian control of hippocampal homeostasis, and the importance of further research into the mechanisms of chronodisruption, particularly during adolescence, to better understand its long-term implications for cognitive function and overall health.

## Introduction

The term ‘chronodisruption’ was introduced to describe ‘a breakdown of phasing internal biological systems appropriately relative to the external’, and it can lead to adverse health effects (Erren et al., 2003; Erren and Reiter, 2009; Tranah et al., 2011). Under normal conditions, the endogenous circadian rhythms synchronize with environmental factors like the day-night cycle, meal timing and social routines. However, the modern society has significantly transformed human lifestyles resulting in persistent desynchronization between internal time and activity patterns (Pilorz et al., 2018).

This social-biological time discrepancy is referred to as Social Jet Lag (SJL) and is operationalized as the difference in sleep midpoint between workdays and free days (Wittmann et al., 2006). Epidemiological data estimates that 70% of the industrialized population experiences SJL (Roenneberg et al., 2012; Rutters et al., 2014) which has been linked to various pathologies (Beauvalet et al., 2017; Girtman et al., 2022; Levandovski et al., 2011). Adolescents and young adults are particularly susceptible (Collado Mateo et al., 2012; van der Vinne et al., 2015) showing an average SJL of 2 hours, 39 minutes(Roenneberg et al., 2012) accompanied by impaired coping, reward function, impulse control and emotional regulation (Claudatos et al., 2019; de Souza and Hidalgo, 2014; Mathew et al., 2019; Nechifor et al., 2020; Sheaves et al., 2016; van der Vinne et al., 2015), which are reflected in decreased academic performance (Díaz-Morales and Escribano, 2015; Haraszti et al., 2014). In fact, the degree of SJL directly correlates with lower educational achievement (Chandrakar, 2017; Smarr and Schirmer, 2018; van der Vinne et al., 2015). Furthermore, SJL causes circadian misalignment (Baron and Reid, 2014) and dysregulation of the molecular circadian clock (Takahashi et al., 2018).

The mammalian molecular clock is an autonomous, intrinsic timekeeping system driven by a network of proteins that maintain oscillatory transcriptional feedback loops (Patke et al., 2020). The core positive limb is composed of the CLOCK:BMAL1 heterodimer, which initiates the transcription of multiple genes (Bunger et al., 2000; Gekakis et al., 1998), including the *period* (*per*) and *cryptochrome* (*cry*) families (Brown et al., 2005; Kume et al., 1999). The resulting PER and CRY proteins conform the negative limb of the loop by inhibiting CLOCK:BMAL1 activity (Chen et al., 2009; Duong et al., 2011; Sangoram et al., 1998). Hence, their degradation terminates the repression phase and restarts the transcription cycle. Additional regulators like NPAS2, DBP as well as nuclear receptors ROR and REV-ERBs, help to fine-tune the core circadian machinery (Akashi and Takumi, 2005; Guillaumond et al., 2005; Preitner et al., 2002; Sato et al., 2004; Triqueneaux et al., 2004). The proper function of this machinery is crucial for regulating the timing of various biological processes throughout the day, thus contributing to homeostatic control. While the daily expression of clock components has been well-characterized, the rhythmic patterns of clock outputs are less understood. It remains unclear if disruption of daily rhythms in clock genes directly translates into a loss of oscillation in output gene expression.

We do know, thanks to animal models, that the molecular clock within the central nervous system (CNS), interacts with critical modulatory systems like the endocannabinoid, dopaminergic, glutamatergic, and GABAergic systems (Cardinali and Golombek, 1998; Chi-Castañeda and Ortega, 2018; Kim et al., 2017; Kim and Reed, 2021; Moore and Speh, 1993; Vaughn et al., 2010). Accordingly, and consistent with homeostatic failure as a basis of pathological conditions, clock malfunctioning leads to neurotransmission dysregulation. Nonetheless, whether this loss of homeostasis stems from disruptions of oscillations in clock-controlled genes and whether this is responsible for the pathological phenotypes remains to be investigated.

Consequently, our understanding of how chronodisruption, specifically SJL, ultimately translates into morbidity is limited. Several protocols have mimicked SJL in adult mice (Haraguchi et al., 2021; Liu et al., 2019; Oneda et al., 2022), reporting memory impairments (Haraguchi et al., 2021). This highlights the relevance of circadian rhythmicity for hippocampal homeostasis, regulating many forms of neuroplasticity (Snider et al., 2018) and supporting memory formation, retrieval and persistence (Eckel-Mahan, 2012; Hasegawa et al., 2019). Nevertheless, preclinical research of adolescent chronodisruption and its impact on cognition is largely neglected despite the higher prevalence of SJL in young humans. Investigating how developmental disruption of circadian rhythms impacts CNS homeostasis is essential to understanding of the pathophysiology of SJL. This study is, to our knowledge, the first systematic attempt to investigate chronodisruption in adolescent mice and its downstream effects on the circadian expression of neurotransmission systems. We first developed a novel chronic circadian disruption protocol during adolescence and characterized the behavioral phenotype. We performed gene expression analyses throughout the 24h cycle to assess daily rhythms in circadian clock genes as well as external targets such as components from the endocannabinoid, glucocorticoid, glutamatergic and GABAergic systems to assess chronodisruption. Overall, our research shows that disruption of adolescent circadian rhythms induces hippocampal memory impairments and a loss of circadian homeostasis in the CNS in adult mice, specifically on the hypothalamus and hippocampus.

## Methods

### Animals

Wild-type male and female C57BL/6 mice at postnatal day 30 (PD30) were purchased from Charles River (Lyon, France) and delivered to our animal facility (UBIOMEX, PRBB). Mice were grouped-housed at a stable temperature (22 °C ± 2) and humidity (55% ± 10%), with food and water *ad libitum*. Animal weights were recorded daily (Supplementary Information, Figure S1). Animal care and experimental protocols were approved by the Animal Ethics Committee (CEEA-PRBB), following European Community Council guidelines (2016/63/EU). All tests were conducted during the dark phase in standard dim light conditions (15-25) lux from PD67 to PD74.

### Youth Jet Lag paradigm

We developed a novel chronic circadian disruption protocol contemplating a short photoperiod combined with a phase delay (Figure1a). The onset of the light phase was labelled as ZT0. The protocol consisted of two different light/dark (LD) conditions. First, an 8L:16D cycle for the first five days (lights ON at ZT0 and OFF at ZT8) shown to induce changes in circadian phasing (Weinert et al., 2005). Then, during the sixth and seventh day we applied a 4h shift on the onset of the light phase (12L:12D, lights OFF at ZT14). On the eighth day, the cycle returned to 8L:16D. Animals subjected to said protocol constituted the experimental group: “youth jet lag” (YJL). The protocol was implemented from PD30 to PD60. When PD60 was reached, all animals spent one week in standard inverted LD conditions (lights ON at ZT0, lights OFF at ZT12) prior to further testing. Control mice were housed under a standard inverted LD cycle throughout the entire procedure.

**Figure 1.**
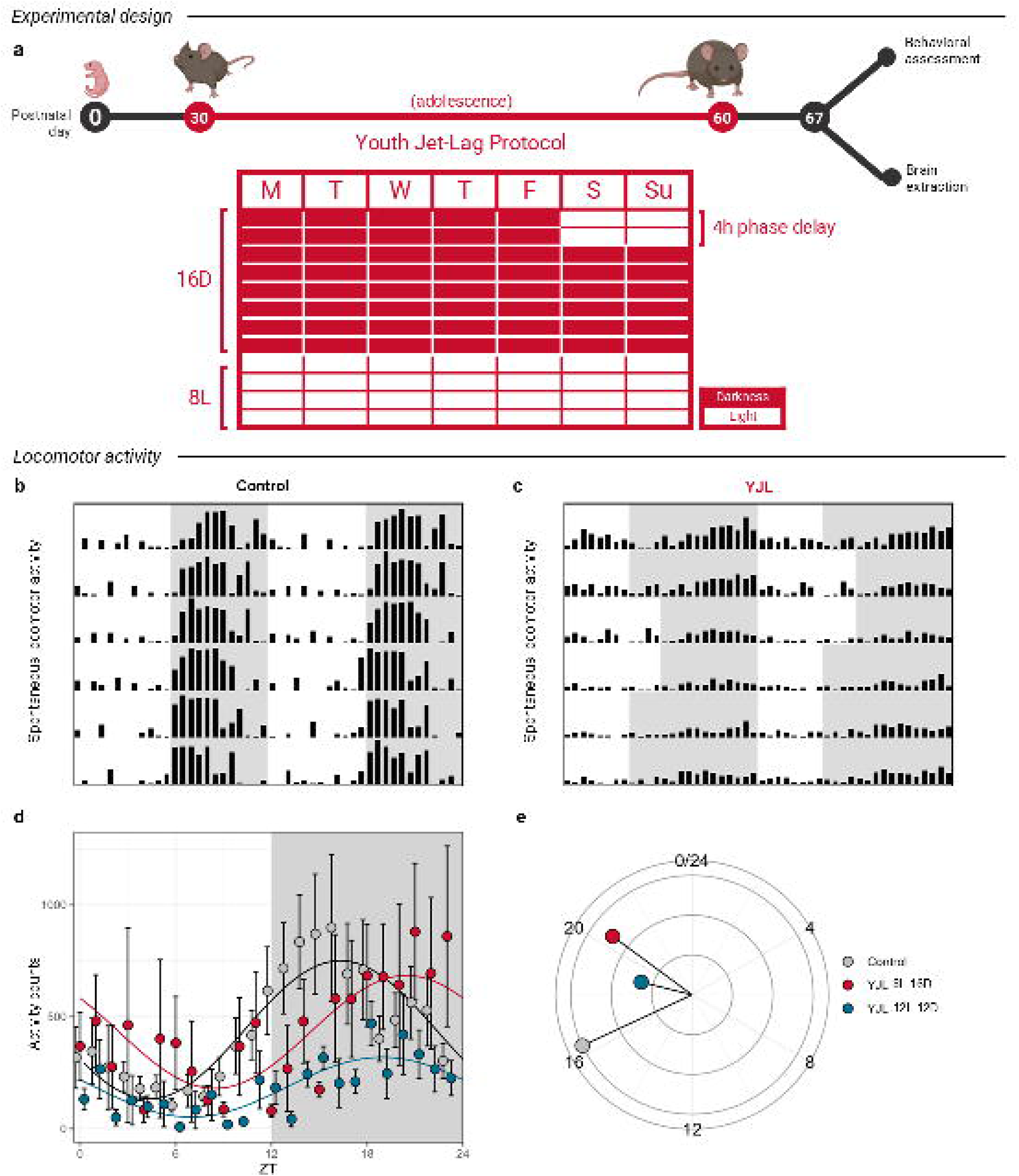
Youth Jet Lag mice display distinct spontaneous locomotor activity patterns during the last week of the protocol. **a** Graphical representation of the experimental design. From PD30 to PD60 animals underwent a disruptive LD cycle with extended activity periods and phase delays. First, a 5-day pattern of 8L:16D, followed by a 4-hour delay in the dark phase onset for 2 days, then returning to 8L:16D. After 5 weeks, at PD60, animals had a one-week under standard LD conditions. Control mice were kept on a consistent 12L:12D cycle throughout the study (PD30 to PD67). Subsequently, animals were used for behavioral experiments (PD67-77) or brain extractions (PD67-68). **b, c** Representative double-plotted actograms from control and YJL. The y-axis shows the monitored days and the x-axis the time of day expressed as ZT. Black bars symbolize spontaneous locomotor activity counts. The darkened areas of the plot represent the ambient illumination regime: grey for lights OFF, white for lights ON. **d** Spontaneous locomotor activity represented as sinusoidal waves representing the least-squares best fit trace for experimental groups; control (grey/black) and YJL activity during the last day of the 8L:16D cycle (red) as well as during the last day of the 12L:12D cycle (blue). Data are expressed as mean (n=4) and standard deviation. **e** Circlegram comparing acrophase and amplitude of spontaneous locomotor activity of the three groups assessed. The lengths of the lines represent the amplitude normalized to control group. The location of the color dots on the circle depicts the ZT of the acrophase. ZT0 represents the onset of the light phase.

### Locomotor Activity

Alteration of daily rhythms in locomotor activity was evaluated by monitoring the locomotion of a small cohort (n = 10) under standard 12L:12D conditions from PD53 to PD60 (LE8825, LE8816; Panlab s.l.u., Barcelona, Spain). For the analyses, we considered the average activity of all control mice at PD57. Alternatively, two distinct days were selected for the YJL group, particularly the last days of the two different light cycles of PD55 (8L:16D) and PD57 (12L:12D).

### Elevated Plus Maze

The elevated plus maze (EPM) (LE-842, Panlab s.l.u., Barcelona, Spain) was used to evaluate anxiety-like behavior, following the methodology of previous studies (Martín-Sánchez et al., 2021) (Figure 2a). Animals’ movement in the maze was assessed by an automated tracking software: Smart (Panlab s.l.u., Barcelona, Spain). The number of entries in each arm was considered and the percentage of time spent in the open arms was calculated with Equation 1.

**Figure 2.**
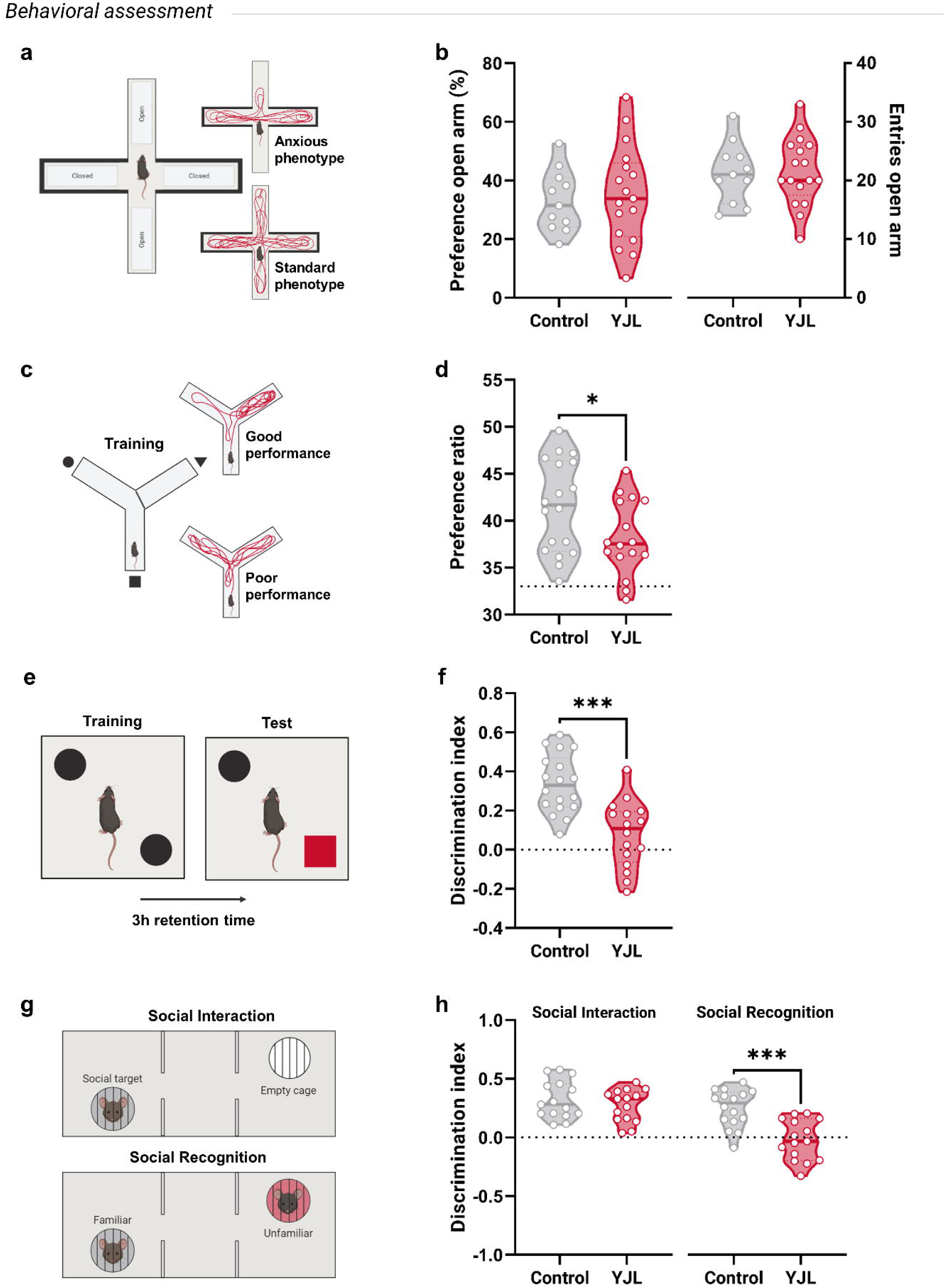
Youth Jet Lag mice display impairments in spatial, short-term, and working social memory but no anxiety-like behavior nor social avoidance. **a** Graphical representation of the EPM apparatus. **b** Percentage of preference (control: n=11; □=33.08% ±10.54, YJL: n=17; x□=35.15% ±16.77) and number of entries in the open arm of the EPM (control: n=11; x□=21.27 ±5.16, YJL: n=17; x□=21.59 ±5.81) **c** Schematic representation of the Reference memory test. **d** Percentage of preference for the novel arm of the Y-Maze in the Reference memory test (control: n=18; x□=41.54% ±4.94, YJL: n=16; x□=38.17% ±3.97). Dotted line at y=33 represents pure chance. **e** Visual representation of the NOR experimental paradigm. **f** Discrimination index during the NOR test (control: n=18; x□=0.336 ±0.15, YJL: n=16; x□=0.079 ±0.17). Dotted line at y=0 represents no preference for either object. **g** Graphical representation of the three-chamber social interaction and recognition test. **h** Discrimination indexes during the three-chamber social interaction (control: n=15; x□=0.323 ±0.16, YJL: n=15; x□=0.273 ±0.14) and social recognition test (control: n=16; x□=0.250 ±0.16, YJL: n=15; x□=-0.021 ±0.17). Dotted line at y=0 represents no preference between interaction cups. Statistical significance was calculated by Student’s *t*-tests. \**p*<.05; \*\**p*<.01; \*\*\**p*<.001. Data are expressed in violin plots as mean, individual values and 25 and 75% percentiles for control (grey) and YJL (red) group.

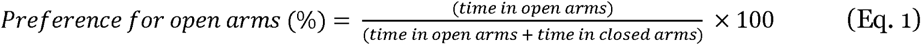

### Reference Memory Test

The reference memory test was performed as previously described (Garcia-Baos et al., 2023) to evaluate spatial reference memory (Figure 2c). The Smart Software tracked the time spent in each of the arms. The preference ratio was then calculated using Equation 2.

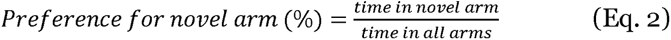

### Novel Object Recognition

The novel object recognition (NOR) test was performed to evaluate short-term declarative memory as described in Garcia-Baos et al., (2023) (Garcia-Baos et al., 2023) (Figure 2e). Both test sessions were video recorded (Logitech C270) and analyzed using the software BORIS (Friard and Gamba, 2016) by a double blinded observer. The discrimination index was computed using Equation 3.

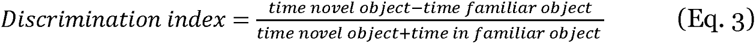

### Social Interaction and Recognition Test

The social interaction and social recognition tests were performed as previously described (Portero-Tresserra et al., 2018) to assess sociability and social memory (Figure 2g). Unfamiliar juvenile mice (PD30) were placed inside a mesh wire cup and used as intruders for the test sessions. The Smart software was used to track the animal and assess the time spent in the areas surrounding the cups. The discrimination index was calculated with Equation 4.

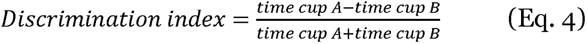

### Time-point tissue collection

Brain sample collection was performed to obtain different measurements of gene expression throughout the 24h cycle to study circadian rhythmicity (Berbegal-Sáez et al., 2024). For this, all animals were housed in a standard LD cycle (12L:12D) from PD60 to PD67. At PD67, mice were sacrificed by cervical dislocation at 6 different time-points: ZT2, ZT6, ZT10, ZT14; ZT18 and ZT22. The hippocampus and hypothalamus regions were isolated (See Supplementary Information), extracted and stored at -80C for further analysis.

### Gene expression analysis

#### RNA isolation and RT-PCR

Total RNA isolation was performed via the TRIZOL method (Supplementary Information). Reverse transcription was performed by High-Capacity cDNA Reverse Transcription Kit (Applied Biosystems, Foster City, CA) following the manufacturer’s protocol, as previously described.

#### OpenArray™ technology

Custom array plates were designed and obtained from Thermo Fisher Scientific Inc. (Supplementary Information, Table S1). To perform the OpenArray™ analyses, 2.5 μl of cDNA sample was mixed with 2.5 μl TaqMan OpenArray™ Real-Time Master Mix (Thermo Fisher #4462159) and loaded into a single well of a 384-well plate (Abraham et al., 2021). Custom OpenArray™ plates were then automatically loaded using the AccuFill System (AccuFill System User Guide, PN4456986) and run in QuantStudio 12K. Data were analyzed using ExpressionSuite Software v1.3 (Thermo Fisher Scientific, 2018-2020). The amplification was normalized to the geometric mean of selected reference endogenous genes *actb*, *gapdh*, *hprt1* and *b2m*. Fold-change values were calculated using the ΔΔCt method using the median value of the ZT2 control as reference sample (Berbegal-Sáez et al., 2024).

### Statistical analysis

We assessed normality and homoscedasticity across all data sets using the Shapiro-Wilk and Spearman’s test. Parametric tests were employed when assumptions of normal distribution and equal variance were met. For single-factor, two-group analyses, we used two-tailed unpaired Student’s *t*-tests. Alternatively, for between-group designs involving two variables, we employed a two-way ANOVA with Bonferroni post-hoc corrections. The present study includes male and female mice in all experiments and considers ‘sex’ as a variable whenever the number of animals was sufficient. When analyses revealed no statistically significant differences based on sex, data were pooled. Statistical significance was set at *p* < .05.

Circadian rhythms were analyzed using the computational tool Kronos (Bastiaanssen et al., 2023; Gheorghe et al., 2024). This software evaluates circadian rhythms within biological datasets by decomposing the time variable into sine and cosine components and using a generalized linear model (GLM) to assess rhythmicity. Sinusoid curves can then be predicted from each outcome variable. The proportion of variance explained by each individual predicted curve with the corresponding *p*-value, along with the acrophase and amplitude is calculated. Additionally, Kronos allows for pairwise comparisons between experimental groups for statistical evaluation of differential rhythmicity.

Lastly, to compare overall gene expression between experimental groups throughout the cycle, we calculated the area under the curve (AUC) and used the total area and standard error to perform unpaired Student’s *t*-tests or with Welch’s correction each gene.

Data were statistically analyzed, and violin plots were generated using GraphPad Prism 8.0. Actograms were created using Microsoft Excel for Microsoft Office 365. Sinusoidal curves and circlegrams were generated using R package Kronos.

## Results

### Youth Jet Lag mice show altered daily rhythms in locomotor activity during the disruptive LD cycle

Spontaneous locomotor activity of YJL and controls was recorded to assess daily rhythms (Figure 1b-e). Sinusoidal curves were generated with Kronos (Figure 1d). The GLM results provided estimated coefficients for the sine and cosine and were found to be significant across all groups (Table 1) with strong evidence of periodicity and varying degrees of goodness-of-fit, as evidenced by the *R*-squared values ranging from .136 to .357 across groups. The acrophase and amplitude for each group is reported in Table 1 and graphically represented as a circlegram in Figure 1e. Furthermore, when assessing differential rhythmicity, the *p*-value for the pairwise models indicated that the rhythmic patterns vary significantly across experimental conditions (*p* < .001). Specifically, the ANOVAs indicated significant differences for ‘control vs YJL 12:12’ (*F*(1, 186) = 52.75, *p* < .0001) and ‘YJL 8:16 and YJL 12:12’ (*F*(1, 186) = 24.29, *p* < .0001). However, no differences were found for ‘control and YJL 8:16’ (Supplementary Information, Table S2).

**Table 1.** Kronos’ rhythmic analysis output of locomotor activity.

| Experimental group | $p$ value | $r$ square | Average | Acrophase | Amplitude | Pairwise models<br>$p$ value |
| --- | --- | --- | --- | --- | --- | --- |
| Control | 1.22E-09 | 0.357 | 440.4 | 16.3 | 308.8 | 0.0004 |
| YJL 8L:16D | 0.001 | 0.136 | 433.7 | 20.5 | 252.4 |  |
| YJL 12L:12D | 4.88E-06 | 0.231 | 186.0 | 18.9 | 133.6 |  |

### Youth Jet Lag mice display impairments in spatial and short-term memory, as well as social recognition but no anxiety-like behavior nor social avoidance

#### Elevated Plus Maze

The results of the EPM test yielded that the means of time spent in open arms were similar: for controls and YJL (Figure 2b). Indeed, no significant differences were found between groups for either the percentage of preference or the number of entries in open arms. The two-way ANOVA showed no significant main effects of ‘sex’ nor interaction ‘sex vs experimental group’ (data not shown).

#### Reference memory

In the reference memory test (Figure 2d) both groups showed a preference for the novel arm. However, the unpaired two tailed *t*-test revealed significant differences between groups (*t*(32) = 2.178, *p* = .0369, *r* = .129), indicating a decreased performance of the YJL group. The two-way ANOVA revealed no significant interaction ‘sex vs experimental group’. Generally, the mean preference of females (41.62%) was higher than the males’ (38.64%) regardless of the experimental group (data not shown). However, this was marginally not significant for the ‘sex’ variable.

#### Novel Object Recognition

During the NOR test (Figure 2f), the average total time of object exploration during the test was 53.99 s (*SEM* = 2.321) and 53.29 s (*SEM* = 2.970) for the controls and the YJL group, respectively, with no significant differences (*t*(34) = 0.1881, *p* = .851, *r* = .001, two-tailed) (data not shown). When assessing the preference for the novel object over the familiar, we found that the control mean discrimination index (0.336, *SEM* = 0.035) indicated a higher preference towards the novel object. Meanwhile, the mean discrimination index of YJL indicated no preference between objects. The unpaired two tailed *t*-test yielded significant differences between groups (*t*(32) = 4.698, *p* < .0001, *r* = .408). The two-way ANOVA revealed no significant main effects of ‘sex’ nor interaction ‘sex vs experimental group’ (data not shown).

#### Social interaction and recognition

Regarding social interaction (Figure 2h, left), the average discrimination indexes for control (0.323, *SEM* = 0.042) and YJL (0.272, *SEM* = 0.036) indicated that both groups showed preference for the intruder mouse over the empty cup. Additionally, the unpaired *t*-test revealed no significant statistical differences in sociability between groups (*t*(28) = 0.919, *p* = .366, *r* = .0293, two-tailed test). Regarding social recognition (Figure 2h, right), the average control discrimination index remained higher than 0 (0.250, *SEM* = 0.040) whereas the mean for YJL plummeted (-0.020, *SEM* = 0.044), indicating no preference between intruders. The unpaired *t*-test revealed significant differences between experimental groups (*t*(29) = 4.546, *p* < .0001, *r* = .414, two-tailed). The two-way ANOVA yielded no significant main effects of ‘sex’ (*F*(1, 27) = 0.465, *p* = .501) nor interaction ‘sex vs experimental group’ (*F*(1, 27) = 0. 001, *p* = .974) for neither evaluated behaviors (data not shown).

### Youth Jet Lag animals show loss of circadian rhythmicity in circadian genes as well as external targets

Regarding circadian clock genes, we found area-dependent significant oscillations (Figure 3 and Supplementary Information, Table S3). In the hypothalamus, the control group showed significant oscillation of *arntl1*, *dbp*, *npas2*, and *per2*, whereas for the YJL group, significant oscillation was reported for *arntl1*, *dbp*, and *npas2*. Although *npas2* was reported to follow a circadian pattern of expression in both groups, the pairwise model indicated that the patterns were significantly different between controls and YJL (*p* = .004). Particularly, we observed an acrophase advancement of 5.6 hours. Meanwhile, in the hippocampus (Figure 3k-t) controls exhibited significant oscillation of *bmal1*, *cry1*, *cry2*, *dbp*, *nr1d2*, *per1* and *per2*, whereas the YJL group showed oscillation in *bmal1*, *cry1*, *dbp*, *nr1d2* and *per2*. Regarding targets outside the molecular clock, Kronos identified significant oscillations in several genes in the hypothalamus (Supplementary Information, Table S4) and the hippocampus (Supplementary Information, Table S5). In the hypothalamus, control animals showed a significant oscillation for: *avp*, *crh*, *crhr1*, *crhr2*, *napepld*, *pparα*, *slc17a6*, and *th* (Figure 4a-h). Alternatively, in YJL, significant oscillations were reported exclusively for *pparα*, and *slc17a6*. While in the hippocampus of controls, we found significant oscillation for *crhr2*, *drd2*, *gabra1*, *gsk3b*, *maoa*, *mgll*, *npy*, *ntrk2*, *slc17a8* and *slc1a2* (Figure 5a-j). In the YJL, however, significant oscillation was only reported for *gsk3b* and *napepld*. Importantly, the pairwise model indicated that although *gsk3b* did oscillate in a circadian manner in the YJL, its sinusoidal curve was significantly different to that of the control (*p* = .04). Particularly, we observed an acrophase advancement of 5.5 hours in the YJL group (Figure 5d).

**Figure 3.**
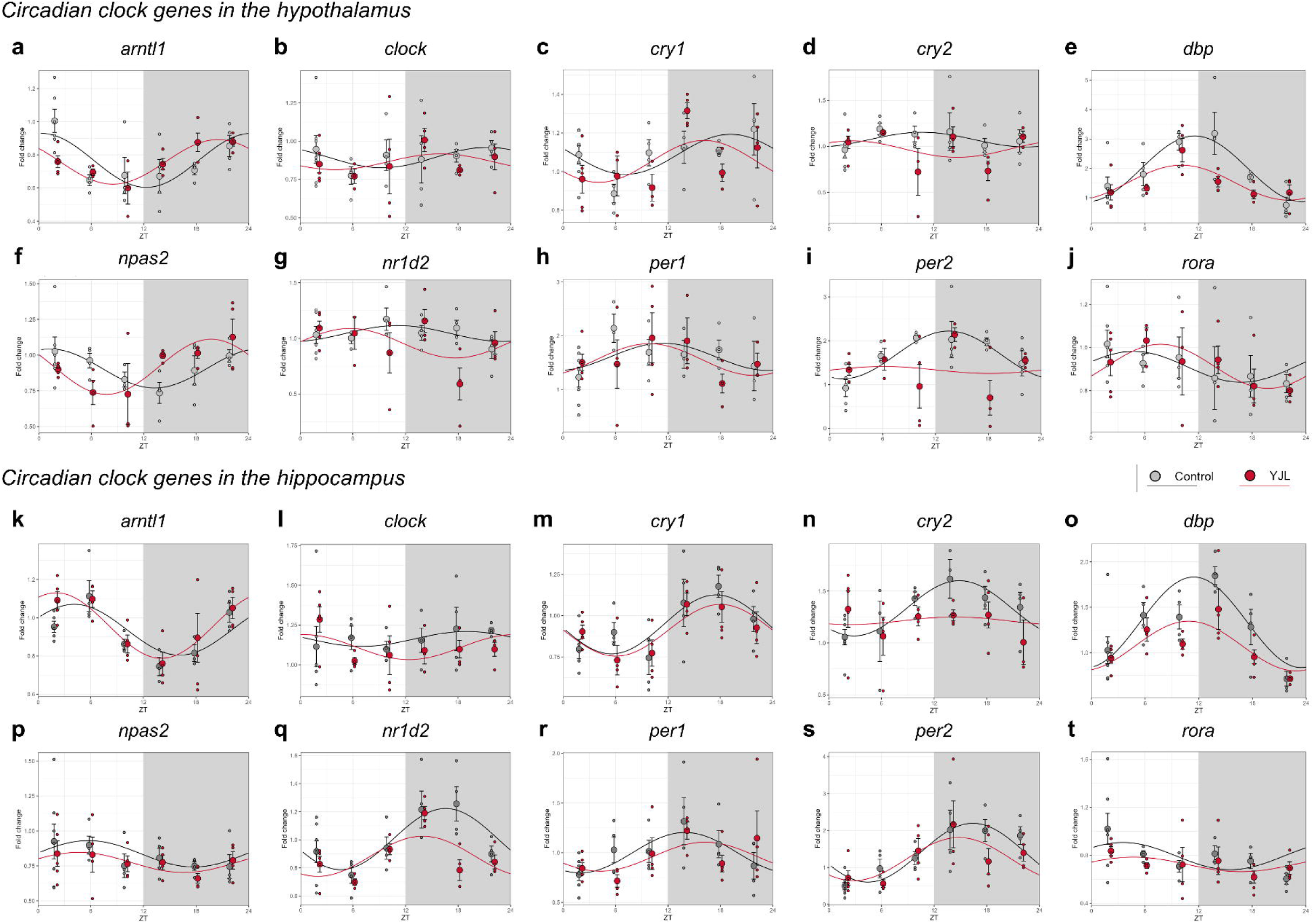
Youth Jet Lag mice display circadian disruption of circadian clock genes in the hypothalamus and hippocampus. Rhythmic analysis output on clock genes mRNA levels in the hypothalamus and hippocampus. Sinusoidal waves fold change of mRNA expression fold change of hypothalamus **a** *artl1*, **b** *clock*, **c** *cry1*, **d** *cry2*, **e** *dbp*, **f** *npas2*, **g** *nr1d2* **h** *per1*, **i** *per2* and **j** *rora*. Sinusoidal waves fold change of mRNA expression fold change of hypothalamus **a** *artl1*, **b** *clock*, **c** *cry1*, **d** *cry2*, **e** *dbp*, **f** *npas2*, **g** *nr1d2* **h** *per1*, **i** *per2* and **j** *rora*. Data is normalized to the median sample of the control group at ZT2. Sinusoid curves represent the least-squares best fit trace for both experimental groups; control (grey/black) and YJL (red). Data are expressed as mean, individual values (n=3-5) and error bars represent standard deviation.

**Figure 4.**
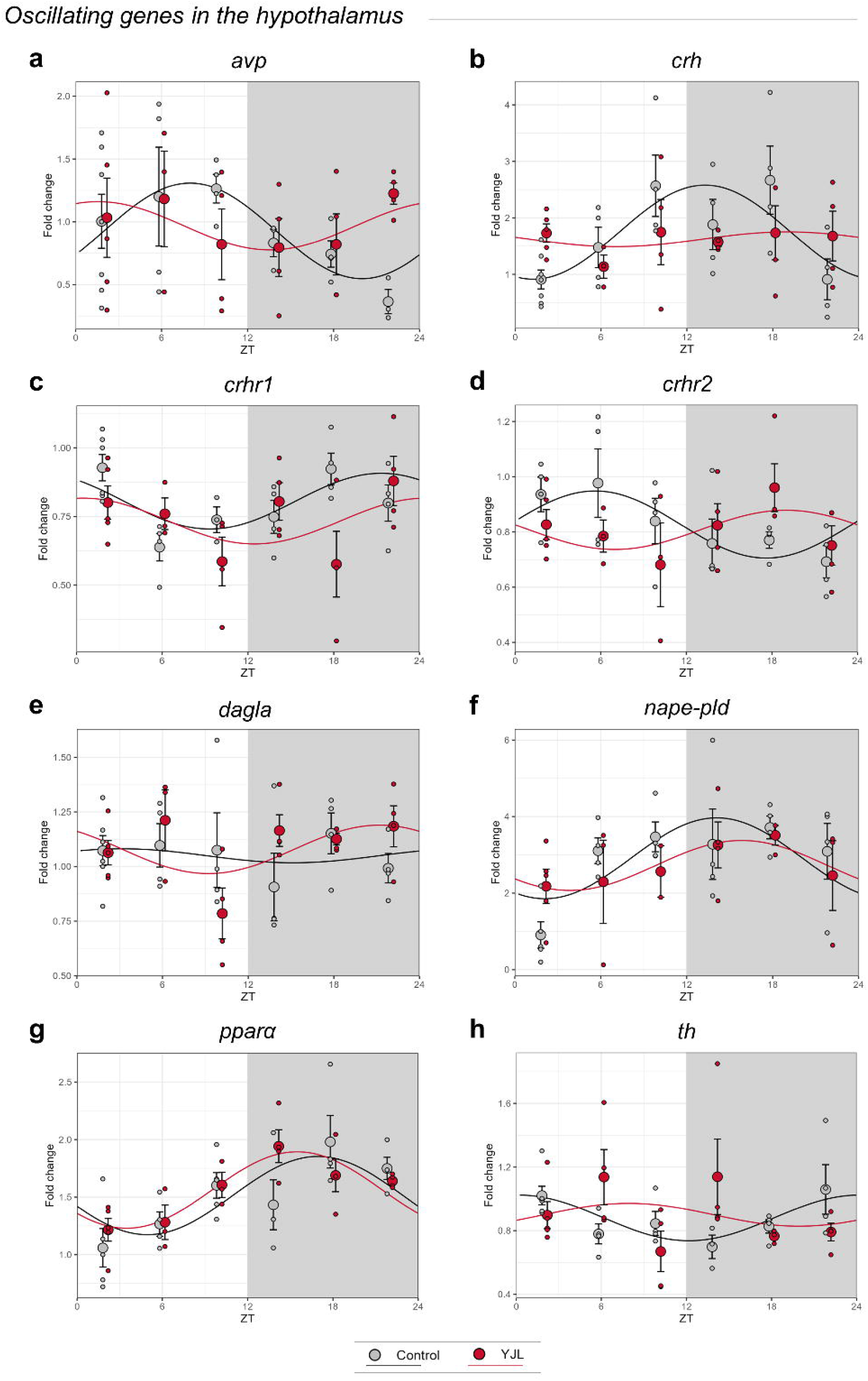
Youth Jet Lag mice display circadian disruption of naturally oscillating genes in the hypothalamus. Rhythmic analysis output on a selection of genes with circadian oscillation levels of mRNA in the hypothalamus. Sinusoidal waves fold change of mRNA expression fold change of **a** *avp*, **b** *crh*, **c** *crhr1*, **d** *crhr2*, **e** *dagla*, **f** *nape-pld*, **g** *pparα* and **h** *th*. Data is normalized to the median sample of the control group at ZT2. Sinusoid curves represent the least-squares best fit trace for both experimental groups; control (grey) and YJL (red). Data are expressed as mean, individual values (n=3-5) and error bars represent standard deviation.

**Figure 5.**
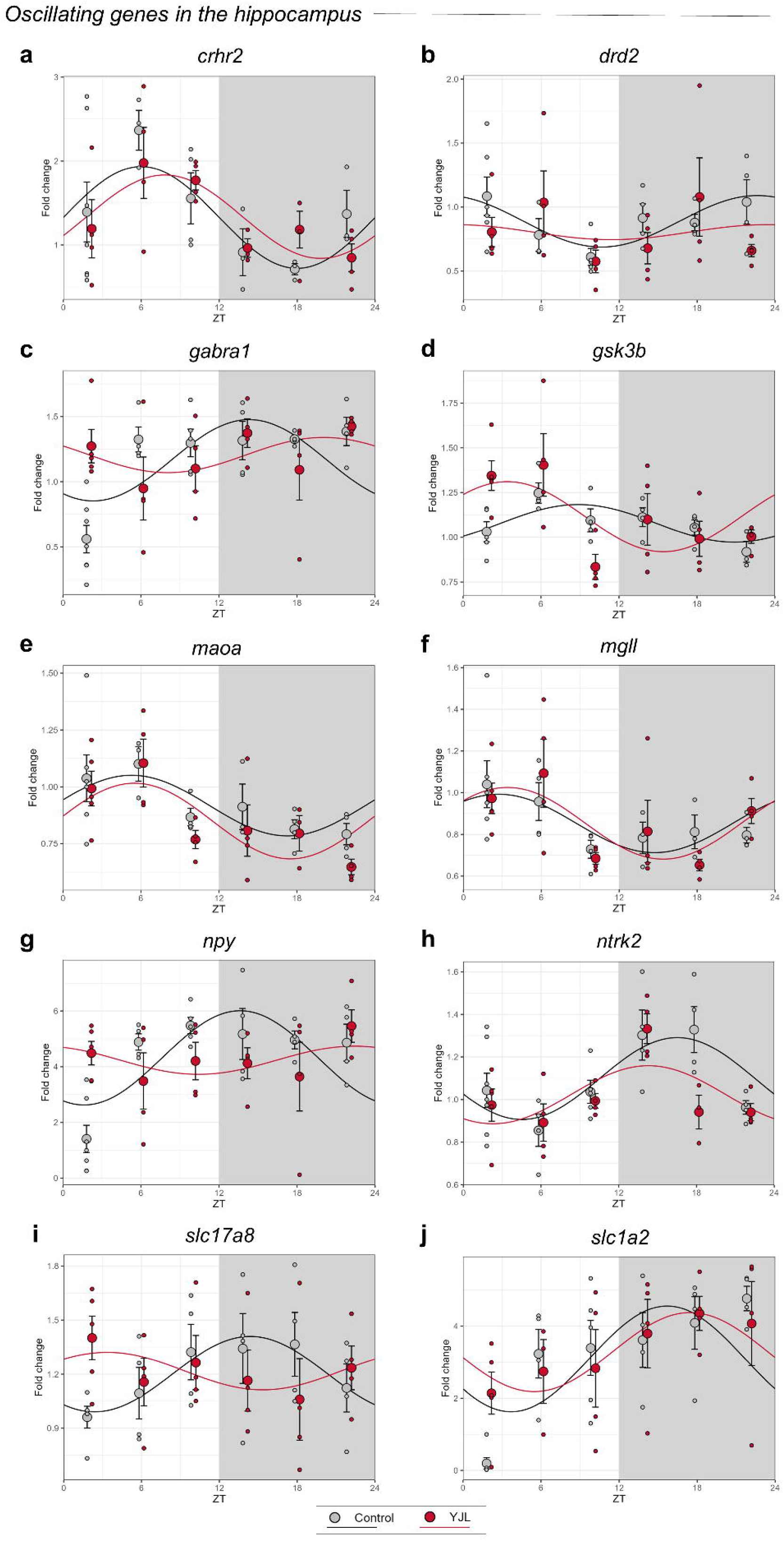
Youth Jet Lag mice display circadian disruption of naturally oscillating genes in the hippocampus. Rhythmic analysis output on a selection of genes with circadian oscillation levels of mRNA in the hypothalamus. Sinusoidal waves fold change of mRNA expression fold change of **a** *crhr2*, **b** *drd2*, **c** *gabra1*, **d** *gsk3b*, **e** *maoa*, **f** *mgll*, **g** *npy*, **h** *ntrk2*, **i** *slc17a8* and **j** *slc1a2*. Data are normalized to the median sample of the control group at ZT2. Sinusoid curves represent the least-squares best fit trace for both experimental groups; control (grey) and YJL (red). Data are expressed as mean, individual values (n=3-5) and error bars represent standard deviation.

## Discussion

The present study provides novel insights into the consequences of chronic adolescent chronodisruption on cognitive function and gene expression rhythms in the CNS. We developed a novel adolescent mouse model of circadian disruption that induces impairments in short-term and spatial memory as well as in social recognition without affecting other cognitive domains like sociability or exploratory and anxiety-like behaviors. Chronodisruption was evidenced by alterations in daily activity rhythms under the disruptive LD cycle, and region-specific changes in clock gene expression. Specifically, in YJL animals, the oscillatory profile of *per2* and *npas2* was altered in the hypothalamus while the hippocampus exhibited a loss of oscillations of *cry2* and *per1*. Furthermore, in basal conditions, we characterized for the first time daily rhythms in the expression of genes such as *crhr1*, *crhr2* in the hypothalamus and *drd2*, *gabra1*, *gsk3b*, *maoa*, *mgll*, *npy*, *ntrk2*, *slc17a8*, *slc1a2* in the hippocampus. Lastly, the YJL group displayed a region-dependent loss of oscillation of gene expression compared to control. On one hand genes related to regulation of the hypothalamic-pituitary-adrenal (HPA) axis and glucocorticoid secretion were affected in the hypothalamus, while those related to neurotransmitter uptake and synaptic plasticity were impacted in the hippocampus. Overall, the present work offers a novel chronodisruption model that recapitulates cognitive deficits observed in adults which suffered SJL during adolescence.

First, LD cycle disruption significantly altered daily locomotor activity rhythms in YJL mice. Control mice exhibited an acrophase consistent with previous reports for the C57BL/6 strain (Dial et al., 2024; Schroeder et al., 2012). However, the YJL group presented a delay in the acrophase during the 8L:16D cycle, a common observation in short photoperiods (Weinert et al., 2005), and indicative of adaptation to the aberrant LD cycle. Additionally, YJL mice showed differential activity patterns relative to the type of cycle they were being exposed to, *i.e.* 8L:16D or 12L:12D with 4h delay of the onset of light phase. These alterations suggest an entrainment to the disruptive LD cycle and point to a potential dysregulation in the timing of circadian processes. Indeed, our molecular analyses of well-reported oscillating clock genes in the hypothalamus, the integrating region of circadian signals, confirm a desynchronization of the circadian machinery evidenced by the loss of daily fluctuations in *per2* and the advancement of *npas2* acrophase. Collectively, these findings provide robust evidence that our model induces chronodisruption.

Regarding behavioral assessment, YJL animals displayed impairments in hippocampal-mediated memory, such as short-term and spatial memory as well as social recognition. This is in accordance with previous studies that have linked circadian disruption to worsened memory performance (Colwell, 2015; Fujioka et al., 2011; Loh et al., 2010). However, existing research typically follows protocols where both chronodisruption and behavioral testing occur during adulthood, rather than developmental periods. Nonetheless, a recent study investigated early-life chronodisruption (from birth until weaning) induced by a chronic advancement of the light phase every other day (Ameen et al., 2022). Behavioral tests were conducted during adulthood and reported impairments in spatial and working memory, aligning with our observations. Additionally, more recently a study has applied a similar approach in order to study the effects of chronodisruption (Bonilla et al., 2024). The authors designed and exposed mice to a protocol with an aberrant LD cycle during adolescence, comparable to our YJL paradigm. The behavioral results reported impairments in the NOR and active avoidance test. These observations are in accordance with the present work, and further provide evidence of the association between adolescent chronodisruption and memory impairments. Hence, this is, to the best of our knowledge, the second study in literature to characterize the effects of chronic circadian rhythm disruption specifically during adolescence. This approach is particularly relevant since during this period, humans exhibit greater susceptibility to circadian disruption due to intrinsically delayed sleep-wake phases and the influence of modern lifestyles.

Although both sexes were included as a variable, it was never found to interact with our experimental manipulation. Unfortunately, contextualizing this finding within existing literature is challenging since most studies on LD-induced chronodisruption are performed exclusively on male mice (Fujioka et al., 2011; Haraguchi et al., 2021; Loh et al., 2010; Oneda et al., 2022). However, and despite known sex differences in circadian biology (Dib et al., 2021), generally, circadian disruption seems to affect male and female rodents similarly (Ameen et al., 2022; Tam et al., 2021). Though, further studies are needed to explore potential sex-specific effects of chronodisruption.

In the context of anxiety-like behavior, exposure to drastically altered LD cycles is considered a stressful experience for the animals (Sakellaris et al., 1975) and chronic stress has been linked to hippocampal memory impairments (Conrad, 2010; Elizalde et al., 2008). Since YJL animals display no increases in anxiety-like behavior, we conclude that the induced memory impairments are unlikely due to stress derived from the exposure to the aberrant LD cycle. In addition, social withdrawal is a hallmark of chronic stress (Tran and Gellner, 2023). Again, since we report no social avoidance, what we observe is a substantially different phenotype to that of anxiety-like behavior arising from a stress-inducing protocol. Hence, the explanation of the worsened memory performance in YJL may be contingent upon chronodisruption.

Indeed, behavioral alterations in YJL were accompanied by molecular evidence of chronodisruption within the hypothalamus, exemplified by alterations in the oscillation of *per2* and *npas2* in YJL. Additionally, in basal conditions, our study is the first in literature to report the oscillatory expression profile of corticotropin-releasing hormone (CRH) receptors *crhr1* and *crhr2* within the hypothalamus. It is known that, in this area, glucocorticoid release follows a strict circadian organization evidenced by daily fluctuations of *crh* mRNA and blood plasma glucocorticoid levels (Kwak et al., 2008). This finding reinforces the control of circadian clock genes in glucocorticoid transmission. In this line, we observe a loss of oscillation of *crh*, *crhr1* and *crhr2* in YJL, suggesting a dysregulation of the HPA axis. Importantly, this can be directly linked to molecular clock alterations, since the function of *per2* in the hypothalamus is tightly related to the control of corticosterone secretion (Russell et al., 2021). Therefore, it is likely that the disruption of daily rhythms of *crh*, *crhr1* and *crhr2* in YJL is associated to the disruption of *per2* oscillations.

Furthermore, both *npas2* and *per2* are key regulators of glutamatergic transmission (Beaulé et al., 2009). Particularly, *per2* is essential for vesicular glutamate transporter 1 (VGLUT1) circadian cycling. Although we did not find significant differences between groups in hypothalamic circadian oscillation in glutamatergic genes (*slc17a6*, *slc17a8* and *scl1a2*), we report a significant overall downregulation in the expression of VGLUT3 (*slc17a8*) in YJL animals (Supplementary Information, Figure S2). Further evidencing the impact of circadian clock genes in hypothalamic neurotransmission.

Since the hippocampus is the main region coordinating various types of memory, molecular analysis was performed in this area. In basal hippocampal conditions, we found circadian oscillation in the expression of core clock genes such as: *bmal1*, *cry1*, *cry2*, *dbp*, *nr1d2*, *per1* and *per2*, which is in accordance with previous literature (Dębski et al., 2020). Furthermore, we describe daily patterns of *crhr2*, *drd2*, *gabra1*, *gsk3b*, *maoa*, *mgll*, *npy*, *ntrk2*, *slc17a8* and *slc1a2* mRNA expression for the first time. These findings highlight the relevance of circadian rhythmicity in hippocampal functioning, controlling relevant processes regarding synaptic plasticity (Eckel-Mahan, 2012).

As expected, adolescent chronodisruption induced a loss in daily expression patterns of clock genes, such as *cry2* and *per1*, as well as those genes whose rhythms of expression have been first characterized by the present study. Interestingly, the alterations in the molecular clock can be linked to the observed memory impairments. Although to date, no role of *cry2* in memory function has been established, it has been observed that reduction of hippocampal *per1* mRNA levels using siRNA prior to training impairs learning, while its overexpression prevents age-related memory deficits (Kwapis et al., 2018). Moreover, PER1-knockout animals perform significantly worse in hippocampal-dependent long-term spatial learning tasks (Jilg et al., 2010). Nonetheless, the specific mechanisms by which *per1* ultimately results in hippocampal memory impairments might be more challenging. It is hypothesized that this effect might occur through the regulation of CREB-mediated transcription since absence of PER1 disrupts both CREB-dependent gene expression and long-term hippocampal memory formation (Rawashdeh et al., 2016, 2014).

In our paradigm, YJL animals show a loss of circadian rhythmicity in a number of genes that are important for memory development. One of the most relevant is *ntrk2*, since TrkB is a well-known potent regulator of hippocampal long-term potentiation (LTP) (Minichiello, 2009). Indeed, in rats, disruptions of the LD cycle impair spatial memory and normal fluctuations in BDNF/TRKB levels (Asadian et al., 2022). Following our observations, additional components that might be contributing to the YJL phenotype are: 1) dopamine receptor D2R which is required for hippocampal-dependent memory and plasticity (Espadas et al., 2020), 2) GluA1 and GluA2 receptors which are determinant contributors to memory retrieval (Pereyra and Medina, 2021), 3) NPY which can exert both inhibitory and stimulatory effects, depending on memory type (Gøtzsche and Woldbye, 2016) or 4) GSK3B, which is required for memory reconsolidation (Kimura et al., 2008). In fact, circadian rhythmicity of GSK3B activation regulates synaptic plasticity in the hippocampus (Besing et al., 2017) and proteasomal degradation of CRY2 which might be related to our observation of altered *cry2* expression. In general, the observed disruption of hippocampal gene expression oscillations may underlie the hippocampal memory impairments exhibited by the YJL animals. This might occur through a *per1* mediated mechanism, although further research is needed to clarify the specific pathway. In line with our findings, a recent study has linked prolonged activation of glucocorticoid receptors to loss of *per1* oscillation in the hippocampus resulting in loss of circadian variation in LTP and impaired memory (Birnie et al., 2023). This observation fits perfectly with our results, in which a loss of hypothalamic *per2* oscillation is accompanied by a dysregulation of *crh*, *crhr1* and *crhr2* gene expression which, in turn, might lead to the disruption of *per1* daily rhythms in the hippocampus potentially resulting in memory impairments. Hence, the interaction between the glucocorticoid system and circadian machinery is a promising mechanism underlying the YJL phenotype.

Finally, some limitations should be considered when interpreting our results. First, the nocturnal nature of mice introduces challenges in extrapolating findings to diurnal species, highlighting the need for caution in generalizing results. Second, circadian rhythm disruption is tightly related to metabolic alterations which could indirectly impact cognition. While weight changes of the animals were monitored and no significant differences were found between groups (Supplementary Information, Figure S1), potential metabolic abnormalities remain a possibility.

Alternatively, a key strength of this work lies in its focus on analyzing variations in gene expression, particularly for targets outside the circadian clock machinery. This approach is crucial since most studies describe either a complete absence or a general reduction/increase in protein and/or mRNA levels. In contrast, our study reveals a loss of circadian oscillation in mRNA levels, without significant changes in overall expression as measured by AUC analysis (Supplementary Information, Figure S2). This finding underscores the limitations of single-time point analysis, since changes in overall expression levels of oscillating molecules could simply reflect a shift in the peak of expression rather than a true up- or down- regulation. Hence, our work emphasizes the importance of incorporating the temporal dimension into molecular analyses. By doing so, we can avoid misinterpreting changes in oscillating components solely based on overall expression levels. This highlights the need for further investigation of the mechanisms underlying the complexity of circadian control of gene and protein expression.

In conclusion, our study provides valuable insights into the complex interplay between circadian rhythms, memory, and daily fluctuations in gene expression. We demonstrated that the YJL model effectively induces chronodisruption, evidenced by altered daily rhythms in locomotor activity and loss of oscillatory patterns in naturally rhythmic genes. Furthermore, we conclude that chronic disruption of circadian rhythms during adolescence induces deficits in short-term, spatial, and social working memory without impacting other cognitive domains such as exploratory behavior, sociability and anxiety-like behavior. Lastly, we report alterations in daily expression of genes associated with glucocorticoid release and synaptic plasticity within the hypothalamus and hippocampus which can be linked to both circadian clock gene disruption and hippocampal memory impairments. This work underscores the critical role of adolescent circadian rhythms in maintaining cognitive function, the relevance of circadian control of hippocampal homeostasis, and the need for future research on the underlying mechanisms by which chronodisruption exerts detrimental health effects.

## Acknowledgements

The authors would like to thank Dr. Thomaz Bastiaanssen and colleagues at the APC Microbiome for their guidance in the fundamentals of the R programming for Kronos analyses and to Dr. Alba García-Baos for her assistance in carrying out the experiments.

This work was supported by the Ministerio de Ciencia e Innovación, “Grant PID2022-136962OB-100 - MCIN/AEI/10.13039/501100011033 and by ERDF A way of making Europe”, Ministerio de Sanidad (Delegación del Gobierno para el Plan Nacional sobre Drogas #2023/005 and #Exp2022/008695 Fondos de Recuperación, Transformación y Resiliencia (PRTR) Unión Europea, and by the Generalitat de Catalunya, AGAUR (#2021SGR00485). IGL received a grant from the Ministerio de Ciencia e Innovación (#PRE2020-091923). PBS received a FI-AGAUR grant from the Generalitat de Catalunya (#2021FI-B00205). OV is recipient of an ICREA Academia Award (Institució Catalana de Recerca i Estudis Avançats, Generalitat de Catalunya).

## Author contribution

IGL and OV were responsible for the study conceptualization. IGL and PBS were responsible for data curation and investigation. IGL was in charge of formal analysis and software. OV was responsible for supervision, project administration and funding acquisition. IGL and OV wrote the original draft. All authors critically reviewed and approved its content.

## Supplementary Material

□ Supplementary Methods
□ Figure S1–S2
□ Table S1–S6

## References

Abraham, N.A., Campbell, A.C., Hirst, W.D., Nezich, C.L., 2021. Optimization of small-scale sample preparation for high-throughput OpenArray analysis. J Biol Methods 8, e143. 10.14440/jbm.2021.339

Akashi, M., Takumi, T., 2005. The orphan nuclear receptor RORalpha regulates circadian transcription of the mammalian core-clock Bmal1. Nat Struct Mol Biol 12, 441–448. 10.1038/nsmb925

Ameen, R.W., Warshawski, A., Fu, L., Antle, M.C., 2022. Early life circadian rhythm disruption in mice alters brain and behavior in adulthood. Sci Rep 12. 10.1038/S41598-022-11335-0

Asadian, N., Parsaie, H., Vafaei, A.A., Dadkhah, M., Omoumi, S., Sedaghat, K., 2022. Chronic light deprivation induces different effects on spatial and fear memory and hippocampal BDNF/TRKB expression during light and dark phases of rat diurnal rhythm. Behavioural Brain Research 418, 113638. 10.1016/j.bbr.2021.113638

Baron, K.G., Reid, K.J., 2014. Circadian misalignment and health. Int Rev Psychiatry 26, 139–154. 10.3109/09540261.2014.911149

Bastiaanssen, T.F.S., Leigh, S.-J., Tofani, G.S.S., Gheorghe, C.E., Clarke, G., Cryan, J.F., 2023. Kronos: A computational tool to facilitate biological rhythmicity analysis. bioRxiv. 10.1101/2023.04.21.537503

Beaulé, C., Swanstrom, A., Leone, M.J., Herzog, E.D., 2009. Circadian Modulation of Gene Expression, but not Glutamate Uptake, in Mouse and Rat Cortical Astrocytes. PLoS One 4, 1–8. 10.1371/journal.pone.0007476

Beauvalet, J.C., Luísa Quiles, C., Alves Braga de Oliveira, M., Vieira Ilgenfritz, C.A., Hidalgo, M.P., Comiran Tonon, A., 2017. Social jetlag in health and behavioral research: a systematic review. ChronoPhysiology and Therapy Volume 7, 19–31. 10.2147/cpt.s108750

Berbegal-Sáez, P., Gallego-Landin, I., Macía, J., Alegre-Zurano, L., Castro-Zavala, A., Welz, P.-S., Benitah, S.A., Valverde, O., 2024. Lack of Bmal1 leads to changes in rhythmicity and impairs motivation towards natural stimuli. Open Biol 14, 240051. 10.1098/rsob.240051

Besing, R.C., Rogers, C.O., Paul, J.R., Hablitz, L.M., Johnson, R.L., McMahon, L.L., Gamble, K.L., 2017. GSK3 activity regulates rhythms in hippocampal clock gene expression and synaptic plasticity. Hippocampus 27, 890–898. 10.1002/hipo.22739

Birnie, M.T., Claydon, M.D.B., Troy, O., Flynn, B.P., Yoshimura, M., Kershaw, Y.M., Zhao, Z., Demski-Allen, R.C.R., Barker, G.R.I., Warburton, E.C., Bortolotto, Z.A., Lightman, S.L., Conway-Campbell, B.L., 2023. Circadian regulation of hippocampal function is disrupted with corticosteroid treatment. Proc Natl Acad Sci U S A 120, e2211996120. 10.1073/pnas.2211996120

Bonilla, P., Shanks, A., Nerella, Y., Porcu, A., 2024. Effects of chronic light cycle disruption during adolescence on circadian clock, neuronal activity rhythms, and behavior in mice. Front Neurosci 18. 10.3389/fnins.2024.1418694

Brown, S.A., Ripperger, J., Kadener, S., Fleury-Olela, F., Vilbois, F., Rosbash, M., Schibler, U., 2005. PERIOD1-associated proteins modulate the negative limb of the mammalian circadian oscillator. Science 308, 693–696. 10.1126/SCIENCE.1107373

Bunger, M.K., Wilsbacher, L.D., Moran, S.M., Clendenin, C., Radcliffe, L.A., Hogenesch, J.B., Simon, M.C., Takahashi, J.S., Bradfield, C.A., 2000. Mop3 Is an Essential Component of the Master Circadian Pacemaker in Mammals. Cell 103, 1009–1017.

Cardinali, D.P., Golombek, D.A., 1998. The Rhythmic GABAergic System. Neurochem Res 23, 607–614. 10.1023/A:1022426519297

Chandrakar, P., 2017. Social jetlag in school students: Evidence to suggest that sleep deprivation during work days is common. Biol Rhythm Res 48, 99–112. 10.1080/09291016.2016.1234026

Chen, R., Schirmer, A., Lee, Y., Lee, H., Kumar, V., Yoo, S.-H., Takahashi, J.S., Lee, C., 2009. Rhythmic PER abundance defines a critical nodal point for negative feedback within the circadian clock mechanism. Mol Cell 36, 417–430. 10.1016/j.molcel.2009.10.012

Chi-Castañeda, D., Ortega, A., 2018. Circadian Regulation of Glutamate Transporters. Front Endocrinol (Lausanne) 9. 10.3389/fendo.2018.00340

Claudatos, S., Baker, F.C., Hasler, B.P., 2019. Relevance of Sleep and Circadian Rhythms to Adolescent Substance Use. Curr Addict Rep 6, 504–513. 10.1007/s40429-019-00277-9

Collado Mateo, M.J., Díaz-Morales, J.F., Escribano Barreno, C., Delgado Prieto, P., Randler, C., 2012. Morningness-eveningness and sleep habits among adolescents: age and gender differences. Psicothema 24, 410–415.

Colwell, C.S., 2015. How a Disrupted Clock may Cause a Decline in Learning and Memory, in: Circadian Medicine. pp. 235–248. 10.1002/9781118467831.ch16

Conrad, C.D., 2010. A critical review of chronic stress effects on spatial learning and memory. Prog Neuropsychopharmacol Biol Psychiatry 34, 742–755. 10.1016/j.pnpbp.2009.11.003

de Souza, C.M., Hidalgo, M.P.L., 2014. Midpoint of sleep on school days is associated with depression among adolescents. Chronobiol Int 31, 199–205. 10.3109/07420528.2013.838575

Dębski, K., Ceglia, N., Ghestem, A., Ivanov, A., Brancati, G., Bröer, S., Bot, A., Müller, J., Becker, A., Löscher, W., Guye, M., Sassone-Corsi, P., Baldi, P., Bernard, C., 2020. The circadian dynamics of the hippocampal transcriptome and proteome is altered in experimental temporal lobe epilepsy. Sci Adv 6. 10.1126/sciadv.aat5979

Dial, M.B., Malek, E.M., Neblina, G.A., Cooper, A.R., Vaslieva, N.I., Frommer, R., Girgis, M., Dawn, B., McGinnis, G.R., 2024. Effects of time-restricted exercise on activity rhythms and exercise-induced adaptations in the heart. Sci Rep 14, 146. 10.1038/s41598-023-50113-4

Díaz-Morales, J.F., Escribano, C., 2015. Social jetlag, academic achievement and cognitive performance: Understanding gender/sex differences. Chronobiol Int 32, 822–831. 10.3109/07420528.2015.1041599

Dib, R., Gervais, N.J., Mongrain, V., 2021. A review of the current state of knowledge on sex differences in sleep and circadian phenotypes in rodents. Neurobiol Sleep Circadian Rhythms 11, 100068. 10.1016/j.nbscr.2021.100068

Duong, H.A., Robles, M.S., Knutti, D., Weitz, C.J., 2011. A molecular mechanism for circadian clock negative feedback. Science 332, 1436–1439. 10.1126/science.1196766

Eckel-Mahan, K., 2012. Circadian Oscillations within the Hippocampus Support Memory Formation and Persistence. Front Mol Neurosci 5. 10.3389/fnmol.2012.00046

Elizalde, N., Gil-Bea, F.J., Ramírez, M.J., Aisa, B., Lasheras, B., Del Rio, J., Tordera, R.M., 2008. Long-lasting behavioral effects and recognition memory deficit induced by chronic mild stress in mice: effect of antidepressant treatment. Psychopharmacology (Berl) 199, 1–14. 10.1007/s00213-007-1035-1

Erren, T.C., Reiter, R.J., 2009. Defining chronodisruption. J Pineal Res 46, 245–247. 10.1111/j.1600-079X.2009.00665.x

Erren, T.C., Reiter, R.J., Piekarski, C., 2003. Light, timing of biological rhythms, and chronodisruption in man. Naturwissenschaften 90, 485–494. 10.1007/S00114-003-0468-6/TABLES/1

Espadas, I., Ortiz, O., García-Sanz, P., Sanz-Magro, A., Alberquilla, S., Solis, O., Delgado-García, J.M., Gruart, A., Moratalla, R., 2020. Dopamine D2R is Required for Hippocampal-dependent Memory and Plasticity at the CA3-CA1 Synapse. Cerebral Cortex 31, 2187–2204. 10.1093/cercor/bhaa354

Friard, O., Gamba, M., 2016. BORIS: a free, versatile open-source event-logging software for video/audio coding and live observations. Methods Ecol Evol 7, 1325–1330. 10.1111/2041-210X.12584

Fujioka, A., Fujioka, T., Tsuruta, R., Izumi, T., Kasaoka, S., Maekawa, T., 2011. Effects of a constant light environment on hippocampal neurogenesis and memory in mice. Neurosci Lett 488, 41–44. 10.1016/j.neulet.2010.11.001

Garcia-Baos, A., Pastor, A., Gallego-Landin, I., de la Torre, R., Sanz, F., Valverde, O., 2023. The role of PPAR-γ in memory deficits induced by prenatal and lactation alcohol exposure in mice. Mol Psychiatry 28, 3373–3383. 10.1038/s41380-023-02191-z

Gekakis, N., Staknis, D., Nguyen, H.B., Davis, F.C., Wilsbacher, L.D., King, D.P., Takahashi, J.S., Weitz, C.J., 1998. Role of the CLOCK protein in the mammalian circadian mechanism. Science 280, 1564–1569. 10.1126/science.280.5369.1564

Gheorghe, C.E., Leigh, S.J., Tofani, G.S.S., Bastiaanssen, T.F.S., Lyte, J.M., Gardellin, E., Govindan, A., Strain, C., Martinez-Herrero, S., Goodson, M.S., Kelley-Loughnane, N., Cryan, J.F., Clarke, G., 2024. The microbiota drives diurnal rhythms in tryptophan metabolism in the stressed gut. Cell Rep 43. 10.1016/J.CELREP.2024.114079/ATTACHMENT/4886320A-F4FA-4E07-9750-0B205FAED8EC/MMC4.XLSX

Girtman, K.L., Baylin, A., O’Brien, L.M., Jansen, E.C., 2022. Later sleep timing and social jetlag are related to increased inflammation in a population with a high proportion of OSA: findings from the Cleveland Family Study. J Clin Sleep Med 18, 2179–2187. 10.5664/jcsm.10078

Gøtzsche, C.R., Woldbye, D.P.D., 2016. The role of NPY in learning and memory. Neuropeptides 55, 79–89. 10.1016/j.npep.2015.09.010

Guillaumond, F., Dardente, H., Giguère, V., Cermakian, N., 2005. Differential control of Bmal1 circadian transcription by REV-ERB and ROR nuclear receptors. J Biol Rhythms 20, 391–403. 10.1177/0748730405277232

Haraguchi, A., Nishimura, Y., Fukuzawa, M., Kikuchi, Y., Tahara, Y., Shibata, S., 2021. Use of a social jetlag-mimicking mouse model to determine the effects of a two-day delayed light- and/or feeding-shift on central and peripheral clock rhythms plus cognitive functioning. Chronobiol Int 38, 426–442. 10.1080/07420528.2020.1858850

Haraszti, R.Á., Ella, K., Gyöngyösi, N., Roenneberg, T., Káldi, K., 2014. Social jetlag negatively correlates with academic performance in undergraduates. Chronobiol Int 31, 603–612. 10.3109/07420528.2013.879164

Hasegawa, S., Fukushima, H., Hosoda, H., Serita, T., Ishikawa, R., Rokukawa, T., Kawahara-Miki, R., Zhang, Y., Ohta, M., Okada, S., Tanimizu, T., Josselyn, S.A., Frankland, P.W., Kida, S., 2019. Hippocampal clock regulates memory retrieval via Dopamine and PKA-induced GluA1 phosphorylation. Nat Commun 10, 5766. 10.1038/s41467-019-13554-y

Jilg, A., Lesny, S., Peruzki, N., Schwegler, H., Selbach, O., Dehghani, F., Stehle, J.H., 2010. Temporal dynamics of mouse hippocampal clock gene expression support memory processing. Hippocampus 20, 377–388. 10.1002/hipo.20637

Kim, J., Jang, S., Choe, H.K., Chung, S., Son, G.H., Kim, K., 2017. Implications of Circadian Rhythm in Dopamine and Mood Regulation. 450 Mol. Cells 40, 450–456. 10.14348/molcells.2017.0065

Kim, R., Reed, M.C., 2021. A mathematical model of circadian rhythms and dopamine. Theor Biol Med Model 18, 1–15. 10.1186/S12976-021-00139-W/FIGURES/6

Kimura, T., Yamashita, S., Nakao, S., Park, J.-M., Murayama, M., Mizoroki, T., Yoshiike, Y., Sahara, N., Takashima, A., 2008. GSK-3beta is required for memory reconsolidation in adult brain. PLoS One 3, e3540. 10.1371/journal.pone.0003540

Kume, K., Zylka, M.J., Sriram, S., Shearman, L.P., Weaver, D.R., Jin, X., Maywood, E.S., Hastings, M.H., Reppert, S.M., 1999. mCRY1 and mCRY2 are essential components of the negative limb of the circadian clock feedback loop. Cell 98, 193–205. 10.1016/S0092-8674(00)81014-4

Kwak, S.P., Morano, M.I., Young, E.A., Watson, S.J., Akil, H., 2008. Diurnal CRH mRNA Rhythm in the Hypothalamus: Decreased Expression in the Evening Is Not Dependent on Endogenous Glucocorticoids. Neuroendocrinology 57, 96–105. 10.1159/000126347

Kwapis, J.L., Alaghband, Y., Kramár, E.A., López, A.J., Vogel Ciernia, A., White, A.O., Shu, G., Rhee, D., Michael, C.M., Montellier, E., Liu, Y., Magnan, C.N., Chen, S., Sassone-Corsi, P., Baldi, P., Matheos, D.P., Wood, M.A., 2018. Epigenetic regulation of the circadian gene Per1 contributes to age-related changes in hippocampal memory. Nat Commun 9, 3323. 10.1038/s41467-018-05868-0

Levandovski, R., Dantas, G., Fernandes, L.C., Caumo, W., Torres, I., Roenneberg, T., Hidalgo, M.P.L., Allebrandt, K.V., 2011. Depression Scores Associate With Chronotype and Social Jetlag in a Rural Population. Chronobiol Int 28, 771–778. 10.3109/07420528.2011.602445

Liu, K., Hou, G., Wang, X., Chen, H., Shi, F., Liu, C., Zhang, X., Han, F., Yang, H., Zhou, N., Ao, L., Liu, J., Cao, J., Chen, Q., 2019. Social Jetlag and Damage to Male Reproductive System: Epidemiological Observation in European and Chinese Populations and Biochemical Analyses in Mice. SSRN Electronic Journal. 10.2139/SSRN.3482809

Loh, D.H., Navarro, J., Hagopian, A., Wang, L.M., Deboer, T., Colwell, C.S., 2010. Rapid Changes in the Light/Dark Cycle Disrupt Memory of Conditioned Fear in Mice. PLoS One 5, 1–12. 10.1371/journal.pone.0012546

Martín-Sánchez, A., Piñero, J., Nonell, L., Arnal, M., Ribe, E.M., Nevado-Holgado, A., Lovestone, S., Sanz, F., Furlong, L.I., Valverde, O., 2021. Comorbidity between Alzheimer’s disease and major depression: a behavioural and transcriptomic characterization study in mice. Alzheimers Res Ther 13, 73. 10.1186/s13195-021-00810-x

Mathew, G.M., Li, X., Hale, L., Chang, A.M., 2019. Sleep duration and social jetlag are independently associated with anxious symptoms in adolescents. Chronobiol Int 36, 461–469. 10.1080/07420528.2018.1509079

Minichiello, L., 2009. TrkB signalling pathways in LTP and learning. Nat Rev Neurosci 10, 850–860. 10.1038/nrn2738

Moore, R.Y., Speh, J.C., 1993. GABA is the principal neurotransmitter of the circadian system. Neurosci Lett 150, 112–116. 10.1016/0304-3940(93)90120-A

Nechifor, R.E., Ciobanu, D., Vonica, C.L., Popita, C., Roman, G., Bala, C., Mocan, A., Inceu, G., Craciun, A., Rusu, A., 2020. Social jetlag and sleep deprivation are associated with altered activity in the reward-related brain areas: an exploratory resting-state fMRI study. Sleep Med 72, 12–19. 10.1016/j.sleep.2020.03.018

Oneda, S., Cao, S., Haraguchi, A., Sasaki, H., Shibata, S., 2022. Wheel-Running Facilitates Phase Advances in Locomotor and Peripheral Circadian Rhythm in Social Jet Lag Model Mice. Front Physiol 13. 10.3389/FPHYS.2022.821199

Patke, A., Young, M.W., Axelrod, S., 2020. Molecular mechanisms and physiological importance of circadian rhythms. Nat Rev Mol Cell Biol 21, 67–84. 10.1038/s41580-019-0179-2

Pereyra, M., Medina, J.H., 2021. AMPA Receptors: A Key Piece in the Puzzle of Memory Retrieval. Front Hum Neurosci 15. 10.3389/fnhum.2021.729051

Pilorz, V., Helfrich-Förster, C., Oster, H., 2018. The role of the circadian clock system in physiology. Pflügers Archiv - European Journal of Physiology 2018 470:2 470, 227–239. 10.1007/S00424-017-2103-Y

Portero-Tresserra, M., Gracia-Rubio, I., Cantacorps, L., Pozo, O.J., Gómez-Gómez, A., Pastor, A., López-Arnau, R., de la Torre, R., Valverde, O., 2018. Maternal separation increases alcohol-drinking behaviour and reduces endocannabinoid levels in the mouse striatum and prefrontal cortex. European Neuropsychopharmacology 28, 499–512. 10.1016/j.euroneuro.2018.02.003

Preitner, N., Damiola, F., Luis-Lopez-Molina, Zakany, J., Duboule, D., Albrecht, U., Schibler, U., 2002. The orphan nuclear receptor REV-ERBα controls circadian transcription within the positive limb of the mammalian circadian oscillator. Cell 110, 251–260. 10.1016/S0092-8674(02)00825-5

Rawashdeh, O., Jilg, A., Jedlicka, P., Slawska, J., Thomas, L., Saade, A., Schwarzacher, S.W., Stehle, J.H., 2014. PERIOD1 coordinates hippocampal rhythms and memory processing with daytime. Hippocampus 24, 712–723. 10.1002/hipo.22262

Rawashdeh, O., Jilg, A., Maronde, E., Fahrenkrug, J., Stehle, J.H., 2016. Period1 gates the circadian modulation of memory-relevant signaling in mouse hippocampus by regulating the nuclear shuttling of the CREB kinase pP90RSK. J Neurochem 138, 731–745. 10.1111/jnc.13689

Roenneberg, T., Allebrandt, K.V., Merrow, M., Vetter, C., 2012. Social Jetlag and Obesity. Current Biology 22, 939–943. 10.1016/j.cub.2012.03.038

Russell, A.L., Miller, L., Yi, H., Keil, R., Handa, R.J., Wu, T.J., 2021. Knockout of the circadian gene, Per2, disrupts corticosterone secretion and results in depressive-like behaviors and deficits in startle responses. BMC Neurosci 22, 5. 10.1186/s12868-020-00607-y

Rutters, F., Lemmens, S.G., Adam, T.C., Bremmer, M.A., Elders, P.J., Nijpels, G., Dekker, J.M., 2014. Is social jetlag associated with an adverse endocrine, behavioral, and cardiovascular risk profile? J Biol Rhythms 29, 377–383. 10.1177/0748730414550199/ASSET/IMAGES/LARGE/10.1177_0748730414550199-FIG1.JPEG

Sakellaris, P.C., Peterson, A., Goodwin, A., Winget, C.M., Vernikos-Danellis, J., 1975. Response of Mice to Repeated Photoperiod Shifts: Susceptibility to Stress and Barbiturates. Proceedings of the Society for Experimental Biology and Medicine 149, 677–680. 10.3181/00379727-149-38877

Sangoram, A.M., Saez, L., Antoch, M.P., Gekakis, N., Staknis, D., Whiteley, A., Fruechte, E.M., Vitaterna, M.H., Shimomura, K., King, D.P., Young, M.W., Weitz, C.J., Takahashi, J.S., 1998. Mammalian circadian autoregulatory loop: a timeless ortholog and mPer1 interact and negatively regulate CLOCK-BMAL1-induced transcription. Neuron 21, 1101–1113. 10.1016/s0896-6273(00)80627-3

Sato, T.K., Panda, S., Miraglia, L.J., Reyes, T.M., Rudic, R.D., McNamara, P., Naik, K.A., Fitzgerald, G.A., Kay, S.A., Hogenesch, J.B., 2004. A functional genomics strategy reveals Rora as a component of the mammalian circadian clock. Neuron 43, 527–537. 10.1016/J.NEURON.2004.07.018

Schroeder, A.M., Truong, D., Loh, D.H., Jordan, M.C., Roos, K.P., Colwell, C.S., 2012. Voluntary scheduled exercise alters diurnal rhythms of behaviour, physiology and gene expression in wild-type and vasoactive intestinal peptide-deficient mice. J Physiol 590, 6213–6226. 10.1113/jphysiol.2012.233676

Sheaves, B., Porcheret, K., Tsanas, A., Espie, C.A., Foster, R.G., Freeman, D., Harrison, P.J., Wulff, K., Goodwin, G.M., 2016. Insomnia, Nightmares, and Chronotype as Markers of Risk for Severe Mental Illness: Results from a Student Population. Sleep 39, 173–181. 10.5665/sleep.5342

Smarr, B.L., Schirmer, A.E., 2018. 3.4 million real-world learning management system logins reveal the majority of students experience social jet lag correlated with decreased performance. Sci Rep 8, 4793. 10.1038/s41598-018-23044-8

Snider, K.H., Sullivan, K.A., Obrietan, K., 2018. Circadian Regulation of Hippocampal-Dependent Memory: Circuits, Synapses, and Molecular Mechanisms. Neural Plast 2018. 10.1155/2018/7292540

Takahashi, M., Tahara, Y., Tsubosaka, M., Fukazawa, M., Ozaki, M., Iwakami, T., Nakaoka, T., Shibata, S., 2018. Chronotype and social jetlag influence human circadian clock gene expression. Sci Rep 8, 10152. 10.1038/s41598-018-28616-2

Tam, S.K.E., Brown, L.A., Wilson, T.S., Tir, S., Fisk, A.S., Pothecary, C.A., van der Vinne, V., Foster, R.G., Vyazovskiy, V. V, Bannerman, D.M., Harrington, M.E., Peirson, S.N., 2021. Dim light in the evening causes coordinated realignment of circadian rhythms, sleep, and short-term memory. Proceedings of the National Academy of Sciences 118, e2101591118. 10.1073/pnas.2101591118

Tran, I., Gellner, A.-K., 2023. Long-term effects of chronic stress models in adult mice. J Neural Transm 130, 1133–1151. 10.1007/s00702-023-02598-6

Tranah, G.J., Blackwell, T., Stone, K.L., Ancoli-Israel, S., Paudel, M.L., Ensrud, K.E., Cauley, J.A., Redline, S., Hillier, T.A., Cummings, S.R., Yaffe, K., for the SOF Research Group, 2011. Circadian activity rhythms and risk of incident dementia and mild cognitive impairment in older women. Ann Neurol 70, 722–732. 10.1002/ana.22468

Triqueneaux, G., Thenot, S., Kakizawa, T., Antoch, M.P., Safi, R., Takahashi, J.S., Delaunay, F., Laudet, V., 2004. The orphan receptor Rev-erbα gene is a target of the circadian clock pacemaker. J Mol Endocrinol 33, 585. 10.1677/JME.1.01554

van der Vinne, V., Zerbini, G., Siersema, A., Pieper, A., Merrow, M., Hut, R.A., Roenneberg, T., Kantermann, T., 2015. Timing of examinations affects school performance differently in early and late chronotypes. J Biol Rhythms 30, 53–60. 10.1177/0748730414564786

Vaughn, L.K., Denning, G., Stuhr, K.L., de Wit, H., Hill, M.N., Hillard, C.J., 2010. Endocannabinoid signalling: has it got rhythm? Br J Pharmacol 160, 530–543. 10.1111/j.1476-5381.2010.00790.x

Weinert, D., Freyberg, S., Touitou, Y., Djeridane, Y., Waterhouse, J.M., 2005. The phasing of circadian rhythms in mice kept under normal or short photoperiods. Physiol Behav 84, 791–798. 10.1016/j.physbeh.2005.03.008

Wittmann, M., Dinich, J., Merrow, M., Roenneberg, T., 2006. Social Jetlag: Misalignment of Biological and Social Time. Chronobiol Int 23, 497–509. 10.1080/07420520500545979

