## Supplementary information for "Chronic circadian disruption in adolescent mice impairs hippocampal memory disrupting gene expression oscillations"

### Supplementary methods

#### Time-point tissue collection

Following cervical dislocation, brains were then removed from the skull and promptly placed on an ice-cold plate, ventral side up. The hypothalamus was first isolated by identification of the Circle of Willis. Then, a curved forceps was used to separate the tissue immediately underlying this anatomical borderline. Subsequently, the brain was rolled over (dorsal side up) to allow hippocampal isolation. For this, a scalpel was used to separate both hemispheres and to remove the white matter from the cortex until the dorsal hippocampi were exposed. Dorsal regions were isolated from adjacent tissues near the third ventricle through angled incisions oriented perpendicular to the longitudinal axis of the hippocampus. The cortex was then gently pushed away with the help of a spatula until reaching the ventral hippocampi. A final cut was then made along delineating them from the surrounding tissues beneath and from the anterior edge of the hippocampus.

#### RNA isolation via TRIZOL method

The RNA extraction method employed TRIZOL reagent for tissue lysis. Each sample received 1 mL of TRIZOL, followed by gentle tissue breakdown using a pipette. After a 5-minute incubation in ice, chloroform was added (200 µL per 1 mL of TRIZOL), softly mixed, and incubated again in ice. Centrifugation at 14,000 rpm for 15 minutes at 4°C facilitated phase separation. The aqueous phase, ideally clear, was then retrieved and added to 500 µL of cold isopropanol (1:2 ratio of TRIZOL). To facilitate the formation of RNA pellets 5 µg of glycogen were added. The mix was then incubated for 30 minutes at -20C, followed by another centrifugation at 14,000 rpm for 15 minutes until visible pellets were formed. The aqueous phase was then eliminated via decantation. Lastly, ethanol (75%) washes were performed thrice. Pellets were air-dried for 30-60 minutes and resuspended in 20 µL of DEPC water. RNA concentration was quantified using a nanodrop spectrophotometer.

### Supplementary Table 1

**Table S1.** Assay IDs and the corresponding targets of the OpenArrayTM Thermo Fisher.

### Supplementary Table 2

|  | Control vs Friday | | | | |
| --- | --- | --- | --- | --- | --- |
|  | Df | SS | Mean Sq | F value | Pr(<F) |
| Cycle | 1 | 4475.67 | 4475.67 | 0.03 | 0.863 |
| Timepoint_cos | 1 | 9036.05 | 9036.05 | 0.061 | 0.806 |
| Timepoint_sin | 1 | 5566715 | 5566715 | 37.34 | 5.68E-09 |
| Cycle:Timepoint_cos | 1 | 1917080 | 1917080 | 12.86 | 0 |
| Cycle:Timepoint_sin | 1 | 142878.5 | 142878.5 | 0.958 | 0.329 |
| Residuals | 186 | 27727217 | 149071.1 |  |  |
|  | Control vs YJL Sunday | | | | |
|  | Df | SS | Mean Sq | F value | Pr(<F) |
| Cycle | 1 | 3148289 | 3148289 | 52.75 | 1.01E-11 |
| Timepoint_cos | 1 | 234551.6 | 234551.6 | 3.93 | 0.049 |
| Timepoint_sin | 1 | 4013708 | 4013708 | 67.25 | 3.83E-14 |
| Cycle:Timepoint_cos | 1 | 648384.2 | 648384.2 | 10.86 | 0.001 |
| Cycle:Timepoint_sin | 1 | 538693.4 | 538693.4 | 9.03 | 0.003 |
| Residuals | 186 | 11101076 | 59683.21 |  |  |
|  | YJL Friday vs YJL Sunday | | | | |
|  | Df | SS | Mean Sq | F value | Pr(<F) |
| Cycle | 1 | 2915356 | 2915356 | 24.29 | 1.83E-06 |
| Timepoint_cos | 1 | 810506.3 | 810506.3 | 6.75 | 0.01 |
| Timepoint_sin | 1 | 2642027 | 2642027 | 22.01 | 5.24E-06 |
| Cycle:Timepoint_cos | 1 | 335662 | 335662 | 2.8 | 0.096 |
| Cycle:Timepoint_sin | 1 | 126710.8 | 126710.8 | 1.06 | 0.306 |
| Residuals | 186 | 22325367 | 120028.9 |  |  |

**Table S2.** Results of pairwise models ANOVA for locomotor activity.

### Supplementary table 3

**Table S3.** Kronos rhythmic analysis output of clock genes in hypothalamus and hippocampus.

### Supplementary Table 4

**Table S4.** Kronos rhythmic analysis output of genes in hypothalamus.

### Supplementary Table 5

**Table S5.** Kronos rhythmic analysis output of genes in hippocampus.

### Supplementary Table 6

**Table S6.** Area under the curve analysis of gene expression throughout the 24h cycle for both the hypothalamus and hippocampus.

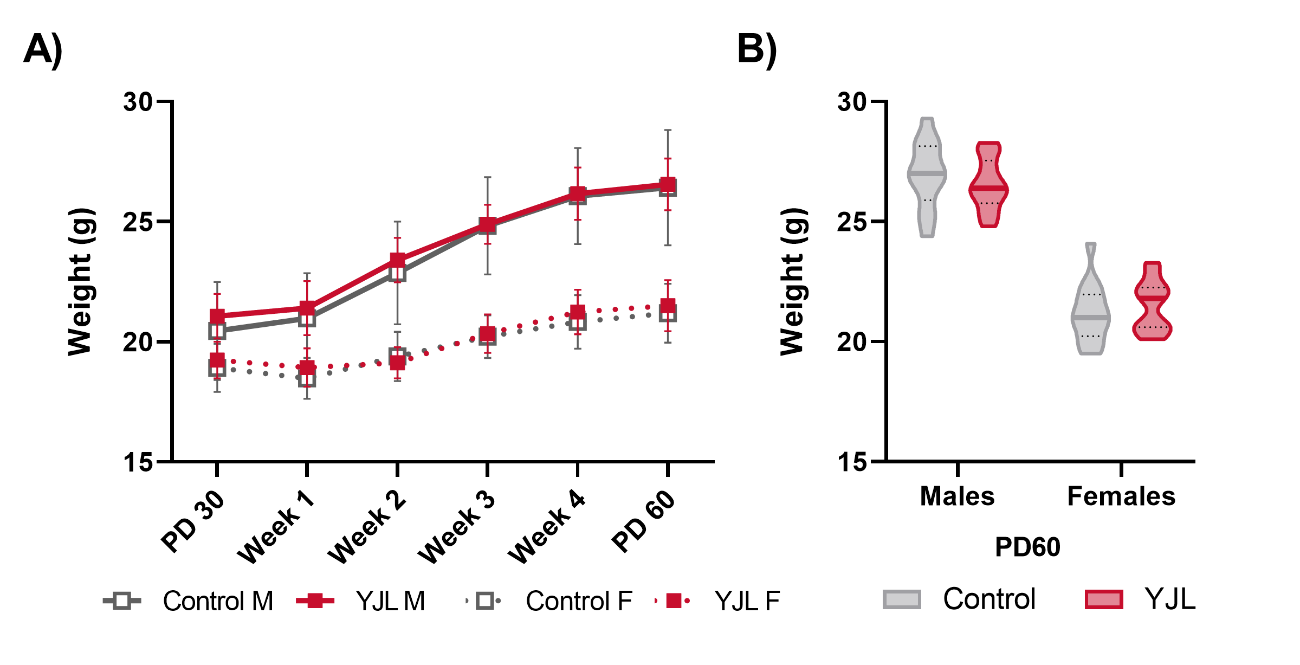

### Supplementary Figure 1

**Figure S1. No significant differences in weight changes between YJL and control during the adolescent-like period.** A) Plot of the animals’ weight at PD30, PD60 and through the duration of the YJL protocol. Statistical significance was calculated with a three-way ANOVA which revealed main effects for ‘time’ (*F*(8, 408) = 167.0, *p* < .0001) and ‘sex’ (*F*(1, 51) = 183.5, *p* < .0001). No significant differences were found between experimental groups (*F*(1, 51) = 0.6252, *p* = .4328). Lastly, significant interactions were observed ‘time vs sex’ (*F*(8, 408) = 29.03, *p* < .0001), while all other interactions were not significant (*p* > .05). Data expressed as mean and standard deviation. B) Significant differences were estimated with a two-way ANOVA which revealed a significant main effect for ‘sex’ (*F*(1, 50) = 273.0, *p* < .0001). No differences were found for ‘experimental group’ (*F*(1, 50) = 0.02185, p = .8831) nor the interaction ‘sex vs experimental group’ (*F*(1, 50) = 1.181, *p* = .2824). Data are expressed in violin plots as mean 25 and 75% percentiles for control (grey) and YJL (red) group.

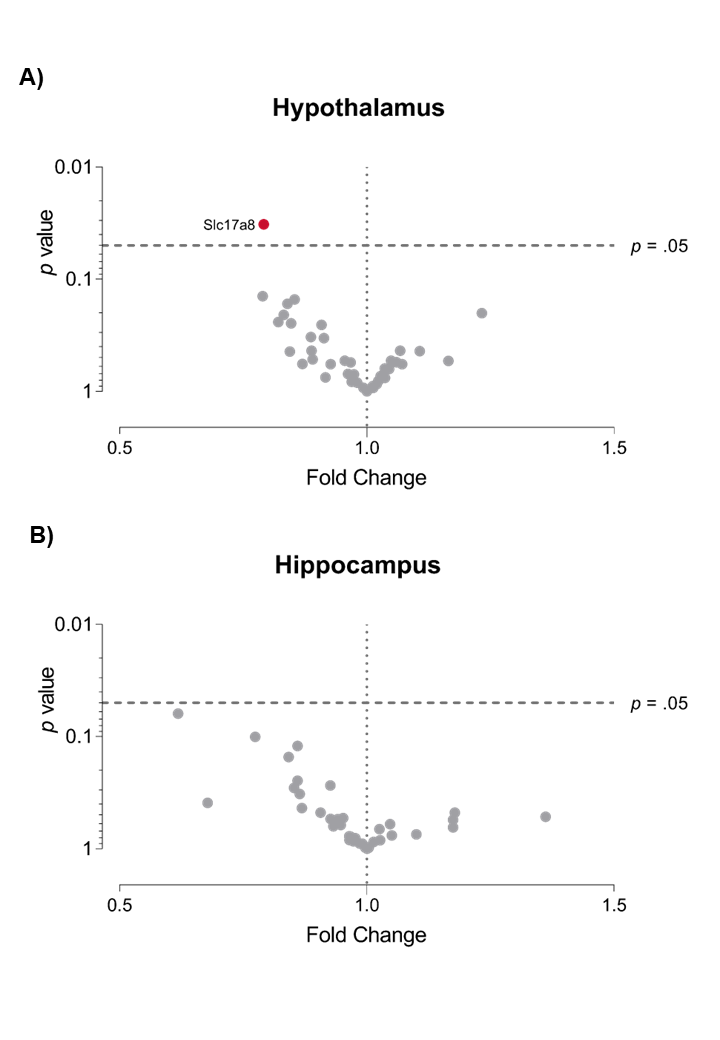

### Supplementary Figure 2

**Figure S2. No differences in total gene expression given by the area under the curve.** Volcano plots showing the results of the differential expression analysis for A) the hypothalamus and B) the hippocampus. Each point represents a gene of interest. The x-axis indicates the fold change in AUC relative to the control. The y-axis represents the -log10 *p*-value of the corresponding unpaired two-tailed Student’s *t*-test with Welch’s correction comparing AUC considering total area and SEMs. The dashed line on the x-axis represents no change in expression (x = 1), while the one on the y-axis indicates the threshold for significance (*p* = .05). Genes that are differentially expressed are shown in red, while those with no significant changes in expression are shown in grey.
